# Asynchronous non-invasive high-speed BCI speller with robust non-control state detection

**DOI:** 10.1101/489013

**Authors:** Sebastian Nagel, Martin Spüler

**Affiliations:** Department of Computer Engineering, Wilhelm-Schickard-Institute for Computer Science, University of Tübingen, 72076 Tüingen

**Author notes:** The authors declare no conflict of interest.

**Keywords:** Electroencephalography (EEG), self-paced, EEG2Code

## Abstract

Brain-Computer Interfaces (BCIs) enable users to control a computer by using pure brain activity. Recent BCIs based on visual evoked potentials (VEPs) have shown to be suitable for high-speed communication. However, all recent high-speed BCIs are synchronous, which means that the system works with fixed time slots so that the user is not able to select a command at his own convenience, which poses a problem in real-world applications. In this paper, we present the first asynchronous high-speed BCI with robust distinction between intentional control (IC) and non-control (NC), with a nearly perfect NC state detection of only 0.075 erroneous classifications per minute. The resulting asynchronous speller achieved an average information transfer rate (ITR) of 122.7 bit/min using a 32 target matrix-keyboard. Since the method is based on random stimulation patterns it allows to use an arbitrary number of targets for any application purpose, which was shown by using an 55 target German QWERTZ-keyboard layout which allowed the participants to write an average of 16.1 (up to 30.7) correct case-sensitive letters per minute. As the presented system is the first asynchronous high-speed BCI speller with a robust non-control state detection, it is an important step for moving BCI applications out of the lab and into real-life.

## Introduction

Brain-Computer Interfaces (BCIs) enable users to control a computer by using brain activity. Their main purpose is to restore several functionalities of motor disabled people, for example, patients who suffered a stroke or have amyotrophic lateral sclerosis (ALS). Probably the most important purpose is to restore the communication ability, therefore, BCI spellers are intensively researched to increase the communication speed. Recent BCI spellers are mainly based on event related potentials (ERPs) or visual evoked potentials (VEPs). The latter are brain responses to visual stimuli and the idea has been proposed by Sutter in 1984 (1), who stated that “the electrical scalp response to a modulated target is largest if the target is located within the central 1° of the visual field” and that “this makes it possible to construct a gaze-controlled keyboard”.

Although recent BCI spellers (2, 3) show high communication speed, they are based on synchronous control, which means that commands are executed in a certain time interval controlled by the BCIs. However, those BCIs are not suitable for real world applications as they cannot differentiate between control and non-control state. Therefore, those BCIs will give a random output if a user is taking a break to think or does not want to control the BCI for other reasons. Therefore, a practical BCI should be asynchronous, or also called self-paced, and should be able to identify the user’s intent to control the system, which is called the “Midas Touch” problem (4). The BCI has to distinguish efficiently between the intentional control (IC) state and the non-control (NC) state, which has been tackled by several methods (5–19).

Some methods make use of hybrid BCIs combining several brain activities, for example, using ERPs (like P300) to distinguish between IC state and NC state in combination with steady-state VEPs (SSVEPs) for classification (6, 12). Others use threshold methods, like Cecotti(15) who developed an asynchronous SSVEP BCI distinguishing between IC state and NC state by normalizing frequency powers for each stimulus frequency following a minimum energy combination approach. Another approach was proposed by Suefusa and Tanaka (14) who developed an asynchronous SSVEP BCI using multiset canonical correlation analysis (MCCA) and a multi-class support vector machine to distinguish between 28 IC classes and the NC class. However, compared to synchronous methods, the current asynchronous BCIs are substantially slower, with 67.7 bit/min being the fastest asynchronous system (14) to date.

The comparison of those methods is difficult, partly because the term “asynchronous” is not uniformly defined, as it is sometimes used for early stopping methods disregarding the NC state detection and sometimes for distinguishing between IC and NC. It also should be mentioned that no unified criteria exist for the evaluation of NC state detection, which makes it difficult to compare the methods between papers. Furthermore, as there are NC state detection methods with fixed trial lengths for synchronous BCIs and approches without fixed trial length for asynchronous BCIs, comparability is even more diffuclt. For the former it is possible to specify the NC detection accuracy and recent works (5–10) achieved accuracies between 76.94 % and 98.91 %. For the latter it is possible to specify the number of erroneous classifications during the NC state per time unit and recent works (11–13) achieved 0.7 to 0.49 erroneous classifications per minute. In summary, all of them did not achieve a reliable NC state detection, as the BCI still executes random commands during the NC state, which decreases the user experience.

In this paper, we present an asynchronous BCI suitable for real-life application. It is an extension of our previous EEG2Code method (20) which predicts the stimulation pattern of an arbitrary VEP response. The method is based on random stimulation patterns which allows to create layouts with an arbitrary number of targets for any application purpose. The presented asynchronous BCI speller achieved a robust NC state detection under different conditions, like reading or looking at another monitor, with only 0.075 erroneous classification per minute. Furthermore, with an average ITR of 122.7 bit/min and up to 205 bit/min it is the first asynchronous high-speed BCI speller.

## Results

### Spelling with an asynchronous EEG2Code BCI

Fig. 1 depicts the components and procedure of the presented asynchronous BCI. It consists of a presentation layer representing a keyboard, an EEG recorder/amplifier, and the asynchronous model that predicts the user-intended target in real-time. The model (EEG2Code) is based on our previous study (20) and is able to predict arbitrary visual stimulation pattern using the spatially filtered EEG. The model prediction is then compared to all possible target patterns to identify the attended target, which is done continuously in an asynchronous fashion. By determining a user-specific threshold, it is detected if the user wants to control the BCI or if the BCI should remain in a non-control state.

**Fig. 1.**
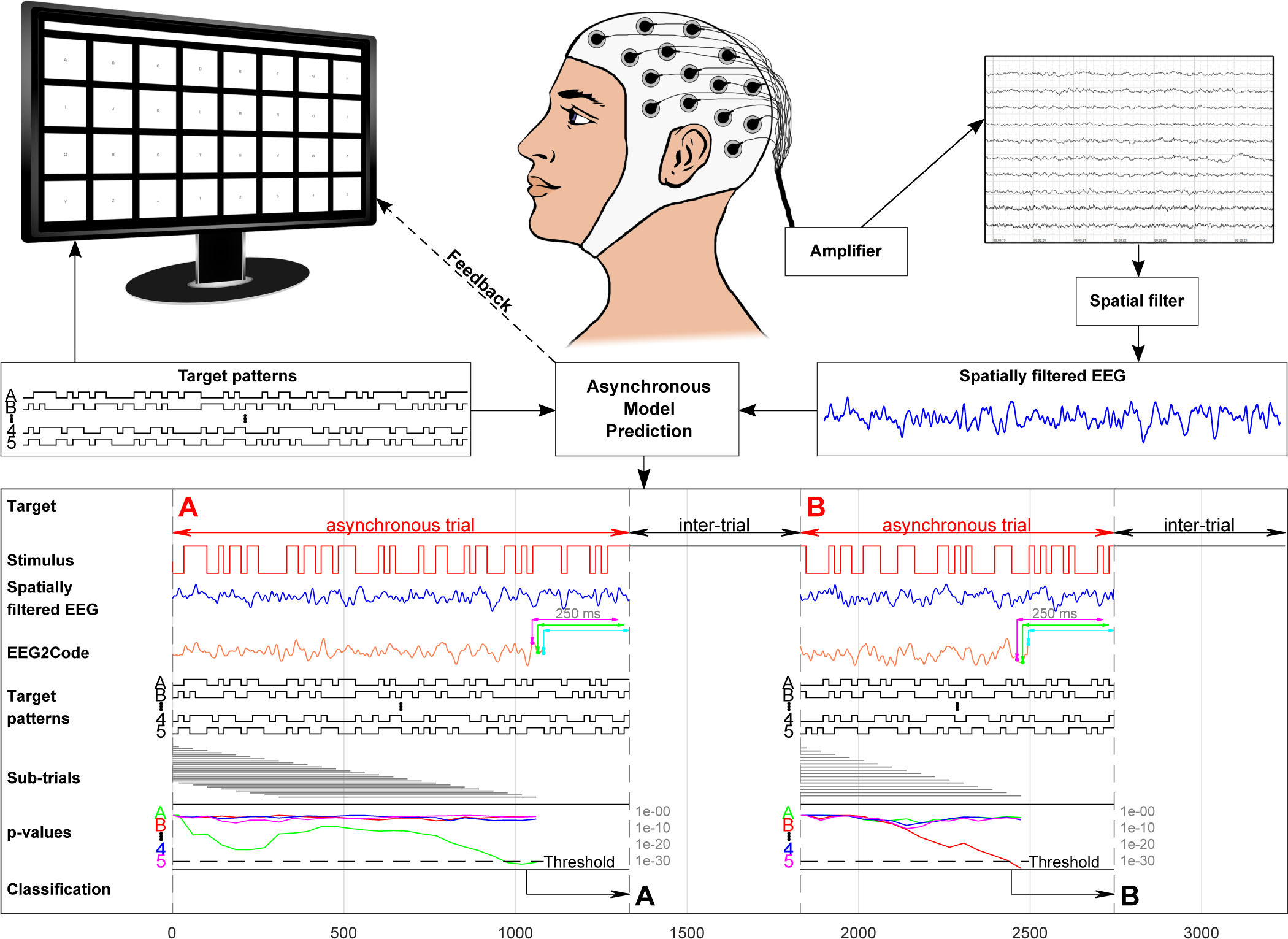
Setup of the asynchronous BCI experiment. The matrix-keyboard layout is as shown on the monitor, it has 32 targets labeled alphabetically from A to Z followed by ‘_’ and numbers 1 to 5. The targets are separated by a blank black space and above targets is the text field showing the written text. Each target is modulated with its own random stimulation pattern. During a trial, the participant has to focus a target. A spatial filter is applied to the measured EEG. For each 250 ms window (slided sample-wise) of the spatially filtered EEG signal (blue line), the EEG2Code model predicts a real value (orange line) which highly correlates to the stimulation pattern. For simplicity, it is only shown for 3 exemplary windows (magenta, green, cyan). Note that the model prediction is delayed by 250 ms because of the sliding window approach. The resulting model prediction is now continuously compared to the stimulation patterns of all targets, not by using the whole trial, but sub-trials (grey lines). We defined, that a sub-trial has a user-specific maximum length (calculated beforehand), once the maximum length is reached, the sub-trial window will be shifted, meaning the beginning of the trial will be discarded. The comparison is done by calculating the p-values with the hypothesis that the correlation coefficient is greater than zero, for simplicity it is only shown for targets A, B, 4 and 5. If at any time one of the p-values is lower than a user-specific threshold (calculated beforehand), the trial stops and the corresponding target will be selected, this is indicated to the participant by highlighting the target in yellow and the corresponding letter is appended to the text field above the keyboard. After an inter-trial time of 0.5 s the next trial starts. Picture modified from (20)

### Online lexicographic spelling performance

For testing the system under optimal conditions, a lexicographic spelling with a 4 × 8 matrix-keyboard (Fig. 1) was used. The left part of Table 1 lists the target prediction accuracy, the average trial time, the corresponding ITR and the corresponding number of correct targets (CT) per minute. The average accuracy was 99.3% ± 0.43% with an average trial duration of 2.61 s ± 0.78 s (including an inter-trial time of 0.5 s), which corresponds to an average ITR of 122.7 bit/min ± 33.2 bit/min and 24.7 CT/min ± 6.8 CT/min, respectively. Across all participants, the minimal and maximal performance ranges from 76.2 bit/min (S08) to 170.9 bit/min (S02).

**Table 1.** Results of the online asynchronous BCI speller. Shown are the results for both the lexicographic spelling (matrix-layout, 32 targets, 192 trials) and the case-sensitive copy-spelling (German QWERTZ-layout, 55 targets, spelling 3 times “Asynchron BCI”). For both the number of correct targets (CT) per minute, the target prediction accuracy, the information transfer rates (ITR) and the average trial duration (including an inter trial time of 0.5 s) are given. Using the matrix-layout, several non-control (NC) states were tested (4 min in total), the results are given as the average number of erroneous classifications per minute during the NC state. Additionally, for the copy-spelling, the number of correct letters (CL) per minute is given, whereas CL takes corrections and case-sensitive letters into account. Best results are in bold font.

| Subject | Matrix-Layout (lexicographic) |  |  |  |  | QWERTZ-Layout (case-sensitive copy-spelling) |  |  |  |  |
| --- | --- | --- | --- | --- | --- | --- | --- | --- | --- | --- |
|  | CT/min | Accuracy [%] | ITR [bit/min] | Time [s] | NC errors/min | CL/min | CT/min | Accuracy [%] | ITR [bit/min] | Time [s] |
| S01 | 25.3 | 99.5 | 125.9 | 2.35 | **0** | 6.2 | 9.5 | 95.2 | 52.4 | 5.99 |
| S02 | **34.6** | 99.0 | **170.9** | **1.72** | **0** | **30.7** | **35.5** | **100.0** | **205.0** | **1.69** |
| S03 | 27.3 | 99.5 | 136.2 | 2.19 | **0** | 21.8 | 25.2 | **100.0** | 145.6 | 2.38 |
| S04 | 15.9 | **100.0** | 80.0 | 3.76 | **0** | 7.6 | 9.8 | 94.3 | 53.2 | 5.80 |
| S05 | 30.5 | 98.4 | 151.1 | 1.94 | 0.25 | 24.3 | 28.0 | **100.0** | 161.9 | 2.14 |
| S06 | 19.7 | 99.5 | 99.5 | 3.02 | **0** | 11.7 | 13.5 | 97.8 | 76.2 | 4.34 |
| S07 | 32.4 | 99.5 | 161.5 | 1.84 | 0.5 | 19.5 | 24.5 | 94.2 | 133.4 | 2.31 |
| S08 | 15.3 | 99.5 | 76.2 | 3.91 | **0** | 4.6 | 5.3 | **100.0** | 30.8 | 11.26 |
| S09 | 19.7 | 99.5 | 99.0 | 3.02 | **0** | 13.6 | 15.7 | **100.0** | 90.9 | 3.81 |
| S10 | 25.7 | 99.0 | 126.9 | 2.31 | **0** | 21.3 | 24.5 | **100.0** | 141.8 | 2.45 |
| mean | 24.7 | 99.3 | 122.7 | 2.61 | 0.075 | 16.1 | 19.2 | 98.2 | 109.1 | 4.22 |

### Online non-control detection performance

As mentioned, the differentiation between IC and NC is important for real-life applications. Therefore, we tested NC states under 4 different conditions, furthermore, both transitions were tested: IC to NC and vice versa. While each IC state was always recognized for all participants, the NC state worked without errors for 8 of the 10 subjects, with 1 error for subject S05 and 2 errors for subject S07. Averaged over all subjects, the NC state detection resulted in 0.075 erroneous classifications per minute (Table 1).

### Online case-senstitive copy-spelling performance

A matrix-keyboard with lexicographic order has the advantage of equal sized targets, but most participants/end-users are familiar with established keyboard layouts. To also evaluate the system under conditions similar to a real-life scenario, we tested the performance using a 55 target German QWERTZ-layout (Fig. S2) by spelling 3 times “Asynchron BCI” (case sensitive). In case of errors, the participants had to correct them by choosing the backspace-target to delete the previous character.

The right part of Table 1 lists the same performance measures as for the matrix-layout, but additionally the number of correct letters (CL) per minute, which includes corrections and case-sensitive letters. The average accuracy was 98.2 % ± 2.56 % with an average trial duration of 4.22 s ± 2.91 s (including an inter-trial time of 0.5 s), which corresponds to an average ITR of 109.1 bit/min ± 56.6 bit/min, 19.2 CT/min ± 9.7 CT/min, and 16.1 CL/min ± 8.7 CL/min, respectively. Interestingly, compared to the matrix layout, for some of the participants the average trial time is highly increased, especially for S01 and S08, which could be explained in part by the reduced target size (5 *×* 5 cm vs. 3 *×* 3 cm).

### Offline threshold optimization

In general, the threshold is used to identify the correct target and to distinguish between IC and NC state. For the online experiment, the threshold was optimized for NC state detection. In order to optimize it for spelling performance, we tested several thresholds by simulating the asynchronous BCI using the lexicographic trials. Fig. 2 shows the accuracies, ITRs, the number of erroneous classifications during NC state and the average trial durations, averaged over all participants. The results show that the communication speed can be optimized to an ITR of 132.4 bit/min with an average trial duration of 1.93 s including 0.5 s inter-trial time. However, using the corresponding threshold during the NC state results in 14.4 erroneous classifications per minute.

**Fig. 2.**
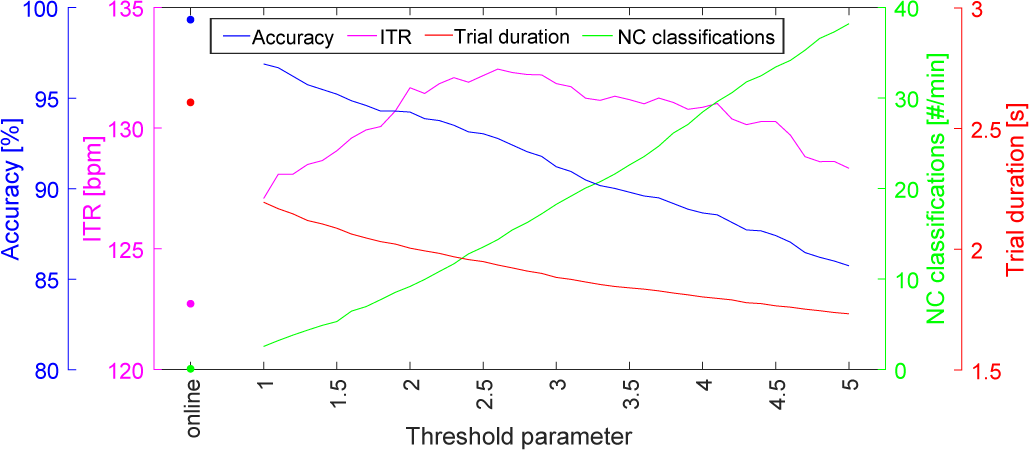
Effects of different p-value thresholds. Shown are the accuracies, information transfer rates (ITRs), erroneous classifications during the non-control state per minute and the average trial duration including an inter-trial time of 0.5 s. The results using the threshold determined during the experiment is marked as “online”. The threshold parameter value defines the percentile of allowed miss-classifications and the resulting threshold is the highest p-value that leads to it (see Methods for details).

## Discussion

For real-life applications, the following factors must be met by a BCI: high communication speed, asynchronous control and non-control state detection. While all three aspects were adressed individually in different publications, the presented system is unique as it combines all three aspects. The presented EEG2Code BCI speller is an asynchronous system that achieves high communication speed (average of 122.7 bit/min), as well as a robust NC state detection with only 0.075 erroneous classifications per minute during the NC state. Compared to the previously fastest asynchronous BCI speller (14) (67.7 bit/min) the ITR is nearly doubled.

As mentioned, among communication speed, an end-user suitable BCI has to distinguish between intentional control (IC) and non-control (NC). Otherwise the BCI will execute random commands during NC states, which highly reduces the user experience. For recent comparable methods for NC state detection (11–13), the best value was 0.49 erroneous classifications per minute during the NC state, which could be reduced by a factor of 6.5 by our method. Here it is worth mentioning that 8 of the 10 participants had a perfect NC state detection without any errors. Since the performance of the NC detection only depends on the threshold, it can easily be optimized for a perfect NC detection. For example, instead of using only a 2 min NC run for threshold determination, the run-time could be increased in order to get the lowest p-value that can occur during a NC state. Indeed, this would reduce the spelling performance by increasing the required trial duration, but this can be counteracted by defining two thresholds: one optimized for IC state and one optimized for NC state. For example, if the user intends to go from IC to NC state a special target has to be gazed, which sets the corresponding threshold to an optimum for NC state. Once the user intends to go back to IC state, the classification of the first intended target takes longer, but afterwards the threshold will automatically be set to the threshold optimized for IC state, which in turn allows to spell faster. We have shown that an optimized threshold results in an increased ITR (132.4 bit/min vs. 122.7 bit/min). Using other methods for threshold optimization could increase the performance even more.

The spelling results revealed high variances between the participants: For example, using the QWERTZ-layout, S02 achieved 205.0 bit/min while S08 achieved only 30.8 bpm. Aside from the physiological difference that can cause such variation, the determination of the optimal maximal sub-trial length is also an important factor. For determination of the optimal sub-trial length we only tested durations between 0.5 s to 3.0 s, but it turned out, that 3.0 s is not enough for participants with a generally bad performance. As shown in Fig. S1, increasing the maximum sub-trial length to 6 s results in a faster classification for S08. Increasing the maximum subtrial length has only one negative effect, it will increase the first classification time after the NC state, but the advantage of faster classifications during the IC state outweighs the disadvantage.

Furthermore, as shown in our previous work (20), the EEG2Code method can be used with an arbitrary number of targets without additional training. We have confirmed that in the present study, although trained using a 32 target matrix-keyboard, the method also works using a 55 target German QWERTZ layout. The results are slightly worse than using the matrix keyboard (109.1 bit/min vs 122.7 bit/min), but this is mainly due to the reduced target size, as this results in lower VEP responses. On the other side, 4 participants achieved an even higher ITR using the QWERTZ-keyboard. Especially, S02 achieved 205.0 bpm resulting in 30.7 correct case-sensitive letters per minute. Furthermore, all participants have noted that a well-known keyboard layout gives a more natural spelling experience, which therefore is another important fact for end-user application.

To summarize, with an average ITR of 122.7 bpm, or an average of 16.1 correctly spelled case-sensitive letters in a realistic scenario, this is the first asynchronous high-speed BCI. It is fully flexible regarding the number of targets and has a near perfect NC state detection. With those properties, the EEG2Code BCI can not only be used for spelling applications, but could also be used for directly controlling mouse and keyboard of the computer (21), and thereby make a huge step towards the application of BCIs in a real-life scenario.

## Materials and Methods

### EEG2Code model

The EEG2Code method was presented in our previous work (20), but for the sake of completeness is described here again, along with the new extensions for asynchronous BCI control and non-control state detection.

### Training

The model is based on a ridge regression model, which uses the EEG signal to predict the stimulation pattern during an arbitrary stimulation. During the experiment, the binary stimulation patterns were presented at a rate of 60 frames per seconds (see section Modulation patterns).

The most prominent parts of a VEP are N1, P1 and N2, the negative/positive potentials with peaks at around 70 ms, 100 ms and 140 ms (post-stimulus), respectively. As the complete VEP lasts for around 250 ms, we use a 250 ms window of spatially filtered EEG data as predictor and the corresponding bit of the stimulation pattern (0=black, 1=white) as response to train the ridge regression model. The window is shifted sample-wise over the data during a trial, meaning that it is required to use 250 ms of EEG data after trial end, otherwise the last 250 ms of a stimulation pattern can not be predicted. Fig. 3 depicts this procedure with a bit-wise window shifting for simplicity. The ridge regression model 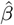 and its bias term *β*_0_ can be calculated by

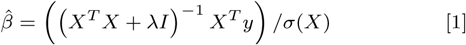

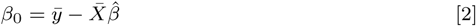

where *X* (the predictor) is a *n* × *k*-matrix with *n* the number of windows and *k* the window length (number of samples). *y* (the responses) is a *n* × 1-vector containing the corresponding bit of the stimulation sequence for each window. *I* is the identity matrix and *λ* the regularization parameter, which was not optimized but set to 0.001. Since a window has a length of *k* = 150 samples, at the used sampling rate *s* = 600*Hz*, the output 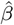 is a coefficient vector of length 150, one for each input sample and the constant bias term *β*_0_. The number of windows *n* depends on the number of trials *N,* the average trial duration *d*, the window length *k* and the sampling rate *s*:

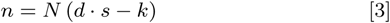

**Fig. 3.**
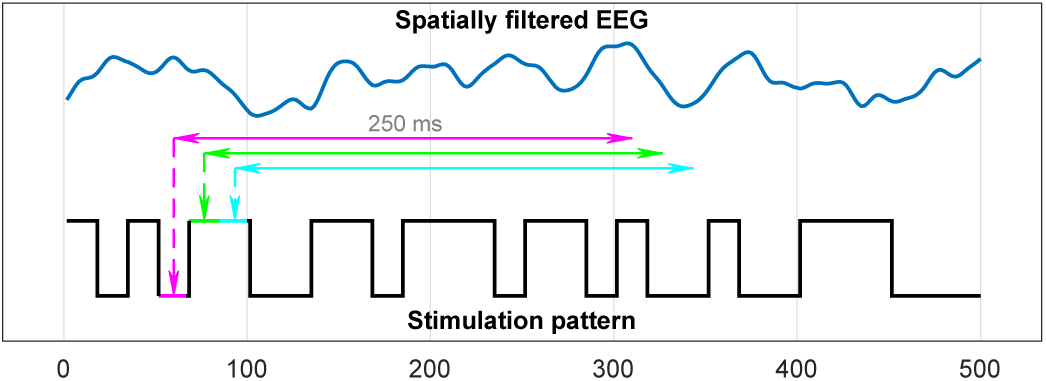
Training of the regression model. Each 250 ms window of the spatially filtered EEG data will be projected to its corresponding bit (1 or 0) of the corresponding stimulation pattern.

As described in section *Data acquisition*, we used *N* = 96 and *d* = 4*s* resulting in *n* = 216, 000 windows used to train the EEG2Code model.

### Prediction

After training, the EEG2Code model is able to predict a sequence of real values, based on the spatially filtered EEG, that highly correlates with the stimulation pattern. The lower part of Fig. 1 depicts the procedure how the EEG2Code model is used to predict a real value *y*_*i*_ for each 250 ms window *i* (sample-wise shifted) of the spatially filtered EEG.

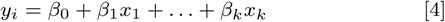

where *β*_0_ is the constant term and *β*_1*…k*_ the coefficients for each window sample *x*_*k*_. The sequence *y* of predicted real values highly correlates with the stimulation pattern. For better understanding, *y* could be transformed to a binary sequence which fits to the binary stimulation pattern with a certain accuracy.

### Asynchronous BCI control

For the asynchronous BCI control, we need a method to choose the correct target out of others based on the EEG2Code model prediction *y* and the classification should only be done if *y* arises from one of the stimulation patterns with a certain probability. As *y* highly correlates with its corresponding stimulation pattern, we calculate the correlation coefficient *r_t_* between the model prediction *y* and the modulation pattern *mt* for each target *t*. The target with the highest *r_t_* should be the correct target. Since the correlation coefficient doesn’t take the length of *y* into account, we calculate the p-value *p_t_* for each target *t* under the hypothesis that the correlation coefficient *r_t_* is greater than zero. The correct target can the be identified by finding the lowest *p_t_* and *p_t_* can also be used as an estimate for the certainty that the selection will be correct.

For better understanding, the complete procedure is depicted in the lower part of Fig. 1. As our approach is not based on a fixed trial length, the p-values are calculated continuously using sub-trial windows. We called those windows sub-trials, on the hand to avoid mix-ups with the windows used for the EEG2Code model prediction (250 ms) and on the other hand to show that the maximum trial length can be longer than the maximum sub-trial length (3 s). If a sub-trial window reaches its maximum length, the sub-trial window will be shifted, which means the beginning of the trial will be discarded. As soon as a p-value *p_t_* is lower than a user-specific threshold (calculated beforehand), the trial stops and the corresponding target *t* will be selected. After an inter-trial time of 0.5 s the next trial starts. If none of the p-values reach the threshold, the trial will continue, which is the case during a non-control phase or if a target can not be classified with a certain probability.

### p-Value threshold and upper sub-trial duration

As different participants do not have exactly the same VEP responses, a user-specific threshold is determined. Furthermore, for better performing participants, shorter sub-trial durations are sufficient, which is why we also determined a user-specific upper sub-trial duration.

We made the assumption that a minimum sub-trial length of 500 ms and a maximum sub-trial length of 3000 ms should be sufficient for all participants. Based on this assumption, we determined the optimal individual sub-trial length for each participant by evaluating different sub-trial length from 500 ms to 3000 ms in 250 ms steps. For each sub-trial length the data was split into corresponding sub-trials and the p-values were calculated as explained above.

We defined the p-value threshold as the p-value of the first percentile out of all sub-trials that lead to a miss-classification. For better understanding, this means that 99 % of all miss-classifications should be excluded by this threshold. The optimal sub-trial length is defined as the shortest duration for which 99 % of all correctly classified sub-trials using that duration have lower p-values as the threshold. If this is not the case for any sub-trial duration, it is set to 3000 ms. This process should find the shortest sub-trial length, for which we can guarantee a IC state detection using the corresponding threshold.

To further optimize the threshold for NC state detection, the participants had to perform a single 2 minute trial where they had to look below the monitor. Using that trial we simulated the asynchronous procedure with the determined upper sub-trial duration. If any p-value occurs which is lower as the determined threshold, it becomes the new threshold. This threshold was then used in the online BCI.

For the offline analysis in the paper, we tested different thresholds. For this, we introduced a threshold parameter, which defines, as explained above, the percentile out of all sub-trials that lead to a miss-classification. The analysis was performed using threshold parameter values between 1 and 5 (in steps of 0.1). The resulting thresholds are no longer optimized for non-control state detection but for optimal spelling performance.

### Modulation patterns

Even thought the EEG2Code model is able to predict arbitrary (random) stimulation patterns, we found that the bit change probability is a crucial property of stimulation patterns that leads to different performances (22). Therefore, we generated a set of 15 bit long (250 ms) sequences with 7 bit changes (50 % bit change probability). This results in a total number of 6,864 bit sequences. As we use a correlation measure to determine the correct target, we generated 100,000 subsets of 150 randomly chosen sequences out of the 6,864 bit sequences and took the subset with lowest average absolute correlation between the sequences in the subset. The resultant subset has an average correlation of -0.004 (SD = 0.276) between any sub-sequence to all others. During the experiment (except for the spatial filter runs), those sequences are randomly assigned to each target. Once the sequence is presented, a new one will be assigned. Therefore, the subset allows to modulate 150^*T/*250ms^ different targets, where *T* denotes the trial duration in milliseconds. The modulation patterns are presented with a rate of 60 bit/s.

### Experimental setup

#### Hardware & Software

The BCI consists of a g.USBamp (g.tec, Austria) EEG amplifier, two personal computers (PCs), Brainproducts Acticap system with 32 channels and a LCD monitor (BenQ XL2430-B) for stimuli presentation. Participants are seated approximately 80 cm in front of the monitor.

PC1 is used for the presentation on the LCD monitor, which is set to refresh rate of 60 Hz and its native resolution of 1920 × 1080 pixels. A stimulus can either be black or white and is synchronized with the refresh rate. The timings of the monitor refresh cycles are synchronized with the EEG amplifier by using the parallel port.

PC2 is used for data acquisition and analysis. BCI2000 (23) is used as a general framework for recording the data of the EEG amplifier and the data processing is done with MATLAB (24). The amplifier sampling rate was set to 600 Hz, resulting in 10 samples per frame/stimulus. Additionally, a TCP network connection was established to PC1 in order to send instructions to the presentation layer and to get the modulation patterns of the presented stimuli. Furthermore, the EEG block-size was set to 32 samples, meaning that a classification is possible each 53,33 ms.

We used a 32 electrodes layout, 30 were located at Fz, T7, C3, Cz, C4, T8, CP3, CPz, CP4, P5, P3, P1, Pz, P2, P4, P6, PO9, PO7, PO3, POz, PO4, PO8, PO10, O1, POO1, POO2, O2, OI1h, OI2h, and Iz. The remaining two electrodes were used for electrooculography (EOG), one between the eyes and one left of the left eye. The ground electrode (GND) was positioned at FCz and reference electrode (REF) at OZ.

#### Presentation layout

We used MATLAB (24) and the Psychtoolbox (25) for the presentation layer. One layout is a 4 × 8 matrix keyboard layout (32 targets in total) as shown on the monitor in Fig. 1, whereas the targets are labeled alphabetically from A to Z followed by ‘_’ and numbers 1 to 5. The targets have a size of 5 × 5 cm are separated by a blank black space and above targets is a text field showing the written text. The second layout is a German QWERTZ-layout, as shown in Fig. S2, including number/symbol row, 2 × shift, caps, tab, backspace, enter, and space (55 targets in total), therefore, it allows to write uppercase and lowercase. The targets with letters/numbers have a size of 3 × 3 cm.

#### Participants

To test the system, 10 healthy subjects were recruited, each participated in one session and completed the whole experiment. All had normal or corrected-to-normal vision and the age ranged from 22 to 34 years. The study was approved by the local ethics committee of the Medical Faculty at the University of Tübingen and conformed to the guidelines of the Declaration of Helsinki. A written informed consent was obtained from all participants.

#### Data acquisition

Initially, the participants performed a run to generate a spatial filter (see *Preprocessing*). The training phase was split into 3 runs (32 trials each) with 4 s trial duration and 1 s inter-trial time, those runs are used for the regression model training and threshold determination. To optimize the threshold for NC state detection, the participants performed a run with a single 2 minute trial where they had to look down (away from the monitor). To test the asynchronous classification, the participants performed 6 runs (32 trials each) with 500 ms inter-trial time. To test the NC state the participants performed 4 runs, each starting with 30 s NC phase followed by 32 IC trials and an additional NC phase with 30 s length. During NC phases, different conditions were tested: monitor was covered, or the participants had to look at the bottom, or they had to look at the left of the monitor, or they had close their eyes. During all mentioned runs the the matrix-keyboard layout was used, 32 trials each in lexicographic order.

Afterwards, each participant had to perform 3 runs using the German QWERTZ-keyboard layout. The participants were asked to write “Asynchron BCI” (case-sensitive) and they should correct any errors that occur. “Asynchron” is the German word for “asynchronous”. It was not prescribed how the participants should spell upper-case or lower-case letters (shift-key or caps-key).

### Preprocessing

#### Frequency filter

The recorded EEG data is bandpass filtered by the amplifier between 0.1 Hz and 60 Hz using a Chebyshev filter of order 8 and an additional 50 Hz notch filter was applied.

#### Correcting raster latencies

Standard computer monitors (CRT, LCD) cause raster latencies because of the line by line image build-up dependent on the refresh rate. As VEPs are affected by these latencies, resulting in a decreased BCI performance, we corrected the raster latencies by shifting the EEG data in respect to its vertical position on the screen (26). For example, with a refresh rate of 60 Hz, the image build-up takes about 16 ms. A target in the (vertical) center of the screen is thereby shown 8 ms after the first pixel is shown, which means that the EEG has to be shifted by 8 ms to correct for that latency. During BCI control, it is not known what character the user wants to select and thereby its position is unknown. Therefore the EEG data is shifted in respect to the target against which it is compared. A more detailed description is given in our previous work (26).

#### Spatial filter

Recent studies (27, 28) have shown increased classification accuracy by using spatial filters to improve the signal-to-noise ratio of the brain signals. As random stimulation is not suitable for spatial filter training, a m-sequence with low auto-correlation is used for target modulation. The training was done using canonical correlation analysis (CCA) as described in a previous work (29), but the presentation layout was as described above and the participants had to perform 32 trials (one per target) whereas one trial lasts for 3.15 s followed by 1.05 s for gaze shifting. As the used m-sequence has a length of 63 bits (1.05 seconds), we got 96 m-sequence cycles per participant, which in turn are used for spatial filter training. The spatial filter is then used for the following experiment.

### Performance evaluation

The BCI control performance is evaluated using the accuracy of correctly predicted targets, the number of correctly predicted targets per minute and the information transfer rates (ITRs) (30). The ITR can be computed with the following equation:

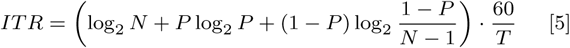

with *N* the number of classes, *P* the accuracy, and *T* the time in seconds required for one prediction. The ITR is given in bits per minute (bit/min). For asynchronous BCI control *N* equals the number of targets (depending on the layout) and *T* the average trial duration including the inter-trial time.

For the NC state, we analyzed the number of erroneous classification per minute during the non-control phases. For the case-sensitive copy-spelling runs (QWERTZ-layout), we also evaluated the number of correct letters which takes corrections and case-sensitive letters into account.

## ACKNOWLEDGMENTS

This work was supported by the *Deutsche Forschungsgemeinschaft* (DFG; grant SP-1533\2-1).

---

**Fig. S1.**
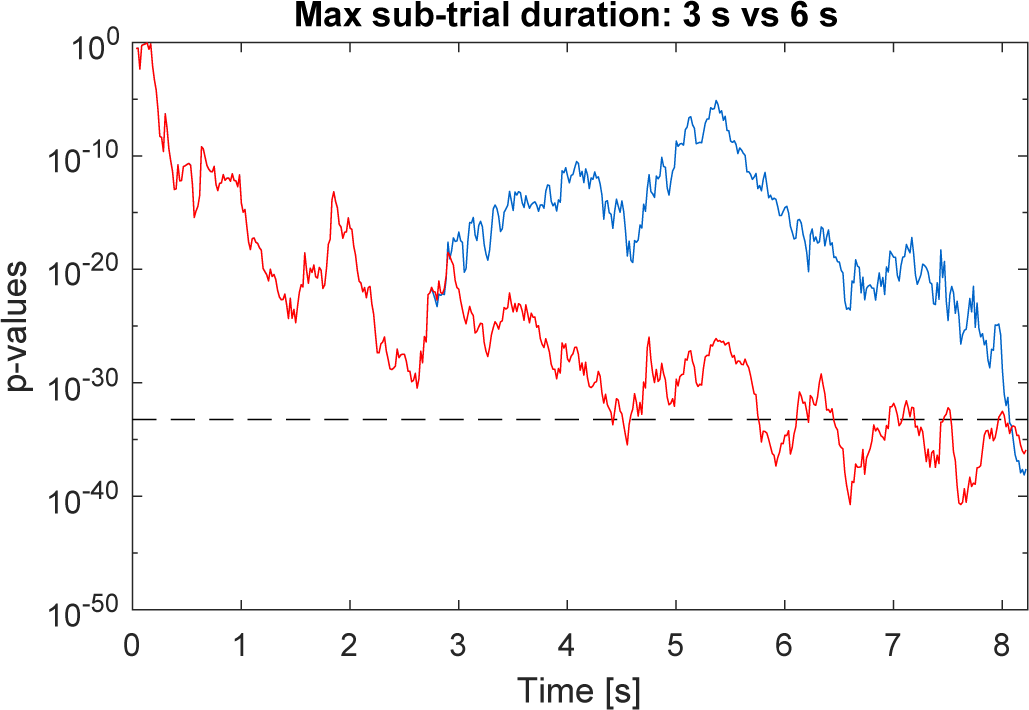
Comparison of the classification speed using different sub-trial durations. The blue line represents the corresponding p-values of the correct target of one of the trials of participant S08 using a sub-trial duration of 3 s which was determined during the online experiment, whereas the red line represents the p-values of the same trial using a sub-trial duration of 6 s. The grey dashed line indicates the p-value threshold used for S08. It clearly shows that using a sub-trial duration of 6 s results in a faster classification speed (approximately 8 s vs. 4.5 s), indicating that a maximum sub-trial length of 3 s is too short for participant S08.

**Fig. S2.**
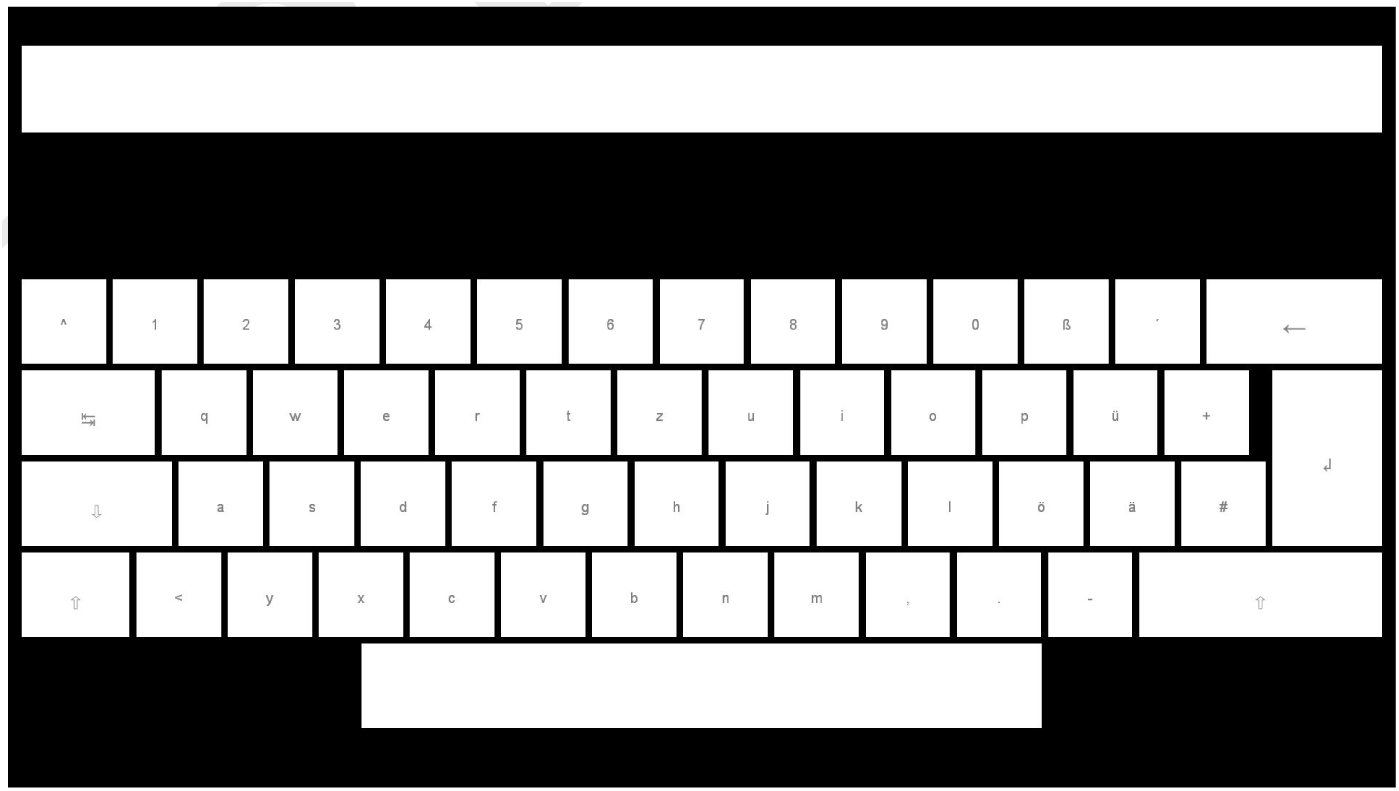
The German QWERTZ-layout, including number/symbol row, 2 *×* shift, caps, tab, backspace, enter, and space (55 targets in total). The targets are separated by a blank black space and above targets is a text field showing the written text.

Author contributions
S.N. developed the methodology, performed the experiment and analyzed the data; S.N. and M.S. designed the experimental procedure and wrote the paper.

